# Distribution of droplets and immune responses after aerosol and intra-nasal delivery of influenza virus to the respiratory tract of pigs

**DOI:** 10.1101/2020.06.04.134098

**Authors:** Veronica Martini, Michael Hinchcliffe, Elaine Blackshaw, Mary Joyce, Adam McNee, Peter Beverley, Alain Townsend, Ronan MacLoughlin, Elma Tchilian

## Abstract

Recent evidence indicates that local immune responses and tissue resident memory T cells (T_RM_) are critical for protection against respiratory infections but there is little information on the contributions of upper and lower respiratory tract (URT and LRT) immunity. To provide a rational basis for designing methods for optimal delivery of vaccines to the respiratory tract in a large animal model, we investigated the distribution of droplets generated by a mucosal atomization device (MAD) and two vibrating mesh nebulizers (VMNs) and the immune responses induced by delivery of influenza virus by MAD in pigs. We showed that droplets containing the drug albuterol, a radiolabel (^99m^Tc-DTPA) or a model influenza virus vaccine (S-FLU) have similar aerosol characteristics. ^99m^Tc-DTPA scintigraphy showed that VMNs deliver droplets with uniform distribution throughout the lungs as well as the URT. Surprisingly MAD administration (1ml/nostril) also delivered a high proportion of the dose to the lungs, albeit concentrated in a small area. After MAD administration of influenza virus, antigen specific T cells were found at high frequency in nasal turbinates, trachea, broncho-alveolar lavage, lungs, tracheobronchial nodes and blood. We conclude that the pig is useful for investigating optimal targeting of vaccines to the respiratory tract.

## Introduction

Respiratory tract infections (RTIs) such as influenza, tuberculosis, respiratory syncytial virus, and recently COVID-19, are a major global health threat, causing considerable morbidity and mortality. Vaccines are the most cost-effective intervention available and much research is targeted towards developing and improving immunization strategies for RTIs. Vaccines are considered effective if they elicit a strong immune response in the peripheral circulation. This systemic cellular and humoral immune response was thought to mediate immunity at sites of disease, whether lung, gut or skin, however, more recent evidence suggests that local immune responses at the site of infection are distinct from systemic responses and play a vital role in protection (1-4). Despite overwhelming evidence highlighting the pivotal role of local immunity, only a handful of mucosal vaccines have been approved for clinical use (5). The major advantage of respiratory delivery of vaccines is that the vaccine is targeted to the site of infection and respiratory immunization efficiently induces lung resident memory (T_RM_) cells, which are important for protective immunity (6-9).

Nevertheless, it is not clear what part of the respiratory tract (RT) should be targeted for optimal protection. Two airway immunization strategies have been developed: local nasal spray and aerosol delivery targeting the lung. In humans, an aerosol measles vaccine has been successfully deployed in Mexico and a live attenuated influenza virus (LAIV) is given to children as a nasal spray (2, 10). However, targeting the lower or upper RT has important safety and immunological implications (11). Studies with measles (12), *Mycoplasma pulmonis* (13), tuberculosis (14) and influenza (15-18) indicate that nasal delivery and lung targeting elicit distinct immune responses. Based on these observations, it is critical to study how vaccines can be optimally delivered to the different areas of the RT in large animal models and humans.

The pig is genetically, immunologically, physiologically and anatomically more similar to humans than small animals (19, 20), making it a useful model for studying the human RT. The lobar and bronchial anatomy of the pig lung is similar to that of humans, they share the same histological structure, epithelial lining, submucosal glands, distribution of sialic acid receptors and electrolyte transport (21). Pigs, like humans, are a natural host for influenza and display similar clinical manifestations and pathogenesis, making them an excellent large animal model for studying influenza infection and new vaccine candidates.

Intra-nasal (i.n.) delivery using a mucosal atomization device (MAD), intra-tracheal instillation and aerosol delivery by nebulizer, have all been used to deliver influenza virus and vaccines to porcine airways (17, 22-26). We have routinely used both MAD and nebulizer for vaccine delivery and viral challenge, but there is very little understanding of where the virus or vaccine is deposited in the porcine RT. Here we evaluated the properties of droplets and devices employed to target the RT in pigs. We compared *in vitro* the droplet size of aerosols generated by a MAD and vibrating mesh nebulizers (VMNs) using the drug albuterol sulphate, a radiolabel (^99m^Tc-DTPA) or the pseudotyped influenza virus vaccine, S-FLU, which has been shown to be protective in mice, ferrets and pigs following RT delivery (24, 25, 27, 28). In this study, the deposition of aerosols generated by the different devices was analyzed *in vitro* using simulated breathing head models *and in vivo* by gamma scintigraphy. Cellular and humoral immune responses after intranasal influenza challenge by MAD were also assessed. These studies will help in the design of rational delivery strategies for vaccines targeting respiratory diseases in livestock species and humans.

## Materials and Methods

### Droplet size evaluation

In vitro experiments were performed with two VMNs producing different droplet size (3 – 5 µm) and the MAD (MAD Nasal™, Wolfe-Tory Medical, US https://www.teleflex.com/usa/en/product-areas/anesthesia/atomization/mad-nasal-device/index.html). Laser diffraction (Spraytec, Malvern Instruments, UK) was used to characterize volumetric median diameter (VMD) of the aerosols generated by the devices (29). Albuterol sulphate (Ventolin, GSK, Ireland) was used as a tracer solution for initial characterization of the VMNs and MAD. The aerosol droplet size of the devices was then characterized using 0.25 ml of technetium (^99m^Tc) complexed with diethylenetriaminepentaacetic acid (DTPA) (^99m^Tc-DTPA) and the candidate influenza vaccine S-FLU vaccine (TCID50 3.5*10^7^/ml). The fine particle fraction and mass median aerodynamic diameter (MMAD) of the droplets were measured by cascade impaction (Next Generation Impactor (NGI), Copley, UK) using 2mg/ml albuterol sulphate. All testing was performed in triplicate.

### *In vitro* albuterol deposition in a pig model

A veterinary mask (Burtons Medical Equipment Ltd, UK) customised with an arrangement of 1-way valves and absolute filters was attached to a 3D printed pig head. The head was connected to a breathing simulator (Dual Phase Control Respirator, Harvard Apparatus, USA) via an absolute filter, onto which inhaled drug was deposited. The breathing simulator was set to reproduce the breathing parameters of a 15 kg (Tidal Volume (Vt) 115 ml, 25 breath per min (BPM), Inhalation:Exhalation (I:E) ratio 1:3) and a 20 kg (Vt 150 ml, 25 BPM, I:E ratio 1:3) pig. A VMN, attached to the mask, was used to nebulise a nominal dose of 1 ml of 2.5mg/2.5ml albuterol sulphate in each test run. The drug eluted from the filter was quantified by UV spectrophotometry (Biochrom UV Vis, Cambridge, UK) at 276 nm and the concentration calculated by interpolation on a standard curve. Results are expressed as a % of the nominal dose (amount in device before administration) delivered.

### *In vitro* lung deposition in an adult human model

An adult head model previously described (29) was attached to the breathing simulator, via an absolute filter, set to simulate a normal adult breathing pattern (Vt 500 ml, 15 BPM, I:E Ratio 1:1). The VMN was connected to an aerosol chamber (Aerogen Ultra, Aerogen, Ireland) and valved aerosol mask (I-Guard, Salter Labs, US) with a supplemental gas flow rate of 2 l/min. Two ml of 1 mg/ml albuterol was delivered. The dose captured on the filter attached at the end of the trachea was calculated as described above.

### *In vivo* pig studies

Two animal studies were performed; each had been approved by the ethical review processes at the respective facility and conducted under valid Procedures Project Licence in accordance with the Animals (Scientific Procedures) Act 1986 amended 2012.

A scintigraphy study was carried out at the University of Nottingham (under PPL 30/3350). Three 6 weeks old Landrace x Hampshire cross, female pigs were obtained from a commercial high health status herd (body weight 10.3 ± 0.25 kg on arrival, 12.6 ± 0.28 kg on pilot leg and 16.9 ±0.34 kg on study leg 3). The animals were sedated with 4.4 mg/kg Zoletil® 100 (Tiletamine with Zolazepam; Virbac, UK) and 0.044 mg/kg Domitor (Medetomidine, Orion Pharma, Finland) prior to administration of radioactive material and remained sedated throughout the imaging period. The experiment was conducted using a randomised crossover design (**Fig. 2A**): the radionuclide ^9m^Tc-DTPA, provided by Radiopharmacy Unit, University of Nottingham, was administered by aerosol using small or medium droplet size VMNs or intranasally with MAD. One ml containing approximately 30 Mega-Becquerel (MBq) of ^99m^Tc-DTPA in 0.9% saline was used for aerosol administration while 10 MBq was delivered intranasally (5 MBq in 1ml per nostril). Experiments were performed at 3-to 4-day intervals to ensure absence of residual radioactivity in the animals and aerosol devices used. Immediately after administration of ^99m^Tc-DTPA, the pigs were imaged under a Mediso X-ring gamma camera fitted with a Low Energy General Purpose collimator (Mediso Medical Imaging Systems, Hungary), and anterior, posterior and lateral images were recorded. A pilot study (**Fig. 2A**) was conducted to ensure optimal dosage and gamma camera settings. Radiolabelled ‘anatomical’ markers were applied around the ears and trunk of pigs for *post hoc* image reconstruction.

**Figure 1.**
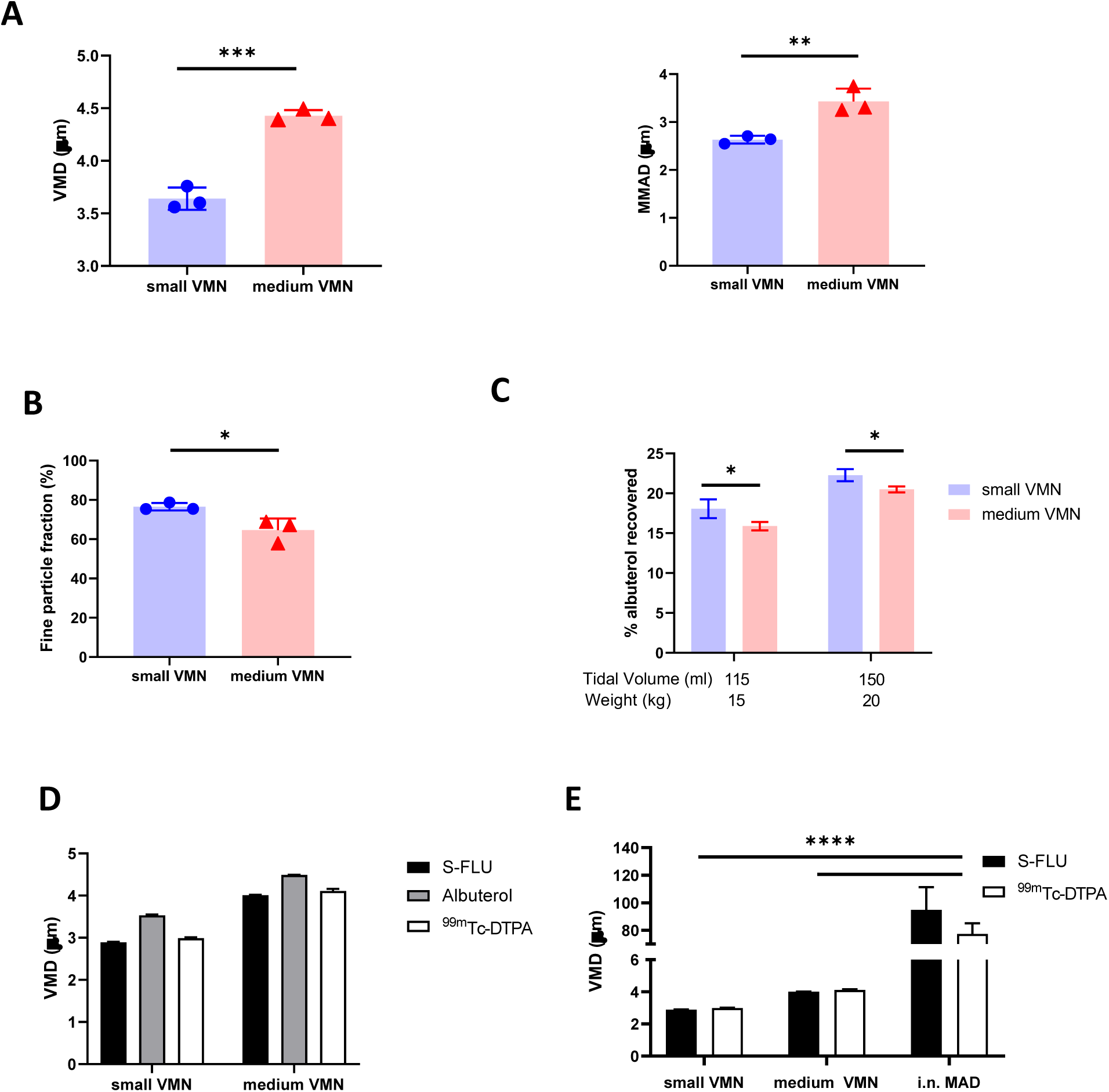
*In vitro* aerosol characterization. **(A)** The volume median diameter (VMD) and mass median aerodynamic diameter (MMAD) were measured for aerosols generated by small droplet size VMN (small VMN) and medium droplet size VMN (medium VMN) using albuterol. **(B)** The fine particle fraction (droplets with diameter smaller than 5 µm) was calculated and compared between the VMNs. **(C)** A 3D printed pig head was used to assess potential lung *in vitro* deposition. Settings of the breathing simulator mimicked a 15 kg pig (115 ml tidal volume (Vt), 25 breaths per min (BPM), Inhale/Exhale ratio (I/E) 1/3) or a 20 kg pig (150 ml Vt, 25 BMP, I/E ratio=1/3). Drug deposition on a filter, interposed between the trachea and breathing simulator, is expressed as a % of the nominal dose (dose in the device before administration). **(D)** The VMDs of aerosols generated using small and medium VMNs with albuterol, S-FLU and ^99m^Tc-DTPA were compared. **(E)** The VMDs of aerosols generated using small and medium VMNs and i.n. MAD with S-FLU and ^99m^Tc-DTPA were analyzed. Unpaired t-test was used for VMD, MMAD, fine particle fraction comparisons. Two-way ANOVA with Tukey post hoc test was used to compare small and medium VMNs and i.n. MAD with S-FLU and ^99m^Tc-DTPA using GraphPad Prism (8.3.0). Two-way ANOVA with Sidak post hoc test was used to compare albuterol aerosol deposition. Asterisks denote *p≤0.05, **p≤0.01, *** p≤0.001 and ****p≤0.0001

**Figure 2.**
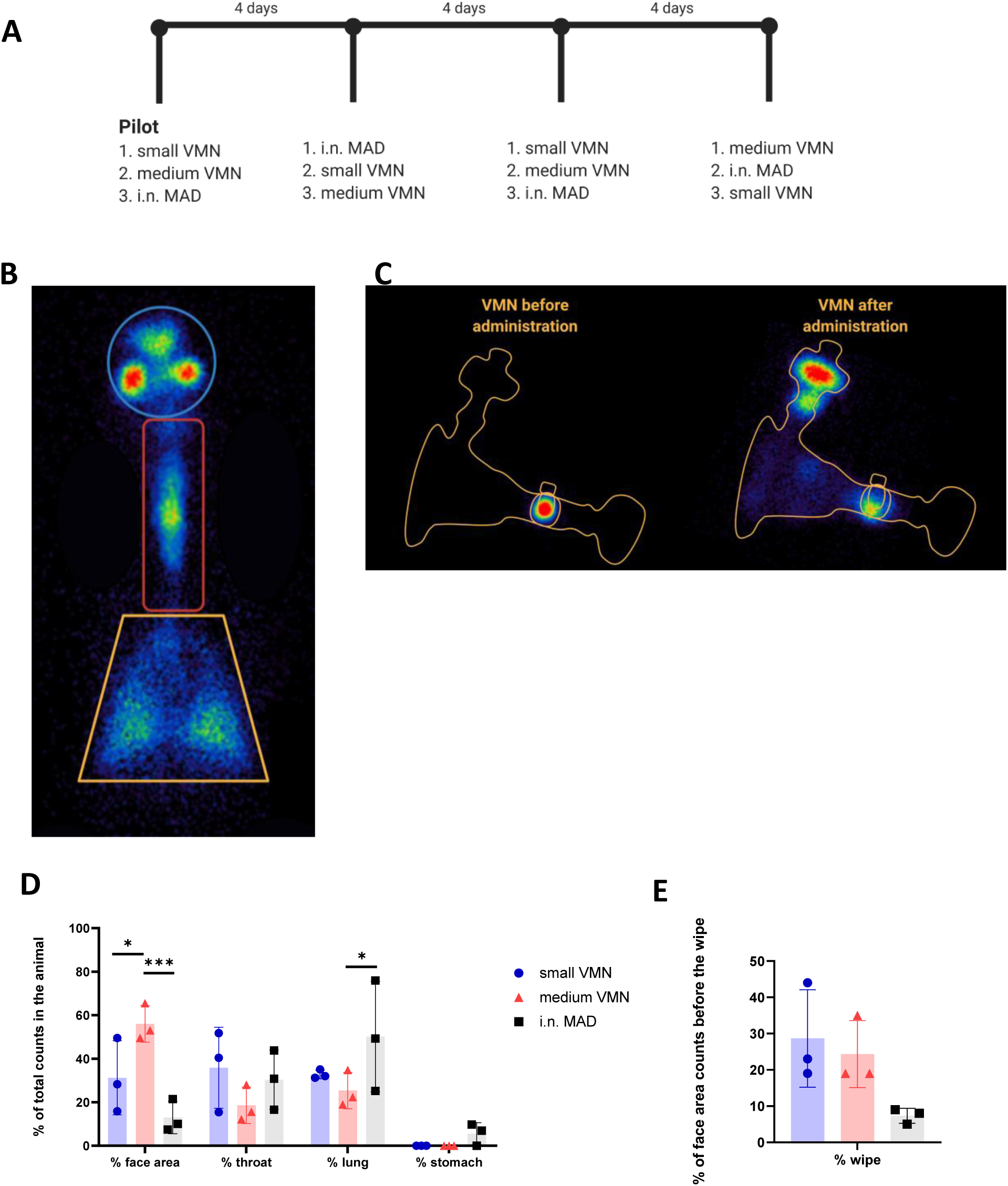
*In vivo* scintigraphy. **(A)** Schematic representation of the cross over experimental design for the *in vivo* scintigraphy study. ^99m^Tc-DTPA was administered to three sedated pigs (indicated with numbers 1,2 and 3) using small and medium VMNs or intranasally with the MAD (i.n. MAD). Each pig received the three different treatments sequentially at 4 day intervals, after the pilot study, conducted to test dose and equipment. **(B)** Images of the VMN before and after delivery. **(C)** Diagram indicating the regions of interest analyzed – face, throat and lung. Radioactive anatomical markers (visible as circles outside the regions of interest) were positioned around the ears and thorax of the pig to enable *post hoc* image reconstruction. **(D)** The counts in each region of interest were expressed as percent of total counts in the animal. **(E)** After recording of all images, the face area of the pigs was wiped. Images of the pig face area and wipe were then recorded. Background corrected wipe counts were expressed as a % of the counts present in the face area before it was wiped. Two-way ANOVA was used for statistical analysis of the deposition in the different region of interest. Kruskal-Wallis test was conducted for the comparison of the face wipes’ radioactivity, using GraphPad Prism (8.3.0) *p≤0.05, and *** p≤0.001.

Influenza challenge was carried out at the Animal and Plant Health Agency (APHA; under PPL P47CE0FF2) by administering 2 ml per nostril of pandemic swine H1N1 isolate, A/swine/England/1353/2009 (pH1N1) at 7.6*10^7^ PFU/ml using MAD to four 9/10 weeks old Babraham inbred pigs. Serum samples and nasal swabs were collected weekly and the pigs were culled 21 days post infection. Peripheral blood mononuclear cell (PBMC), broncho-alveolar lavage (BAL), lung tissue (a sample of every lobe), trachea, nasal turbinate and tracheobronchial lymph nodes (TBLN) were collected at post mortem, mononuclear cells were isolated and cryo-preserved as previously described(25).

### Image analysis

Image analysis was performed using Hermes software (Hermes Medical Solutions, UK). Various anatomical regions of interest (ROIs) were defined on the scintigraphic images: the face area (nasal cavities and external skin) was set as a circle of 10 cm diameter, the throat as a rectangle of length 17 cm and a lung trapezoid was drawn in order to cover the whole lung area (**Fig. 2B**). Radioactive material deposited in the stomach was as a rectangle below the lung area. Counts in the individual ROI were then adjusted for background and decay relative to first image taken for each animal. Deposition in each specific ROI was calculated relative to total counts as follows:

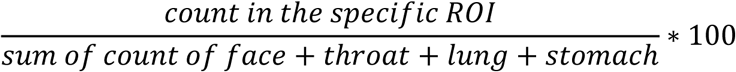

An outline representation of a pig was fitted to the scintigrams to aid visualisation of deposition patterns and it is acknowledged that this might not provide a totally accurate anatomical representation (**Fig. 3A**).

**Figure 3.**
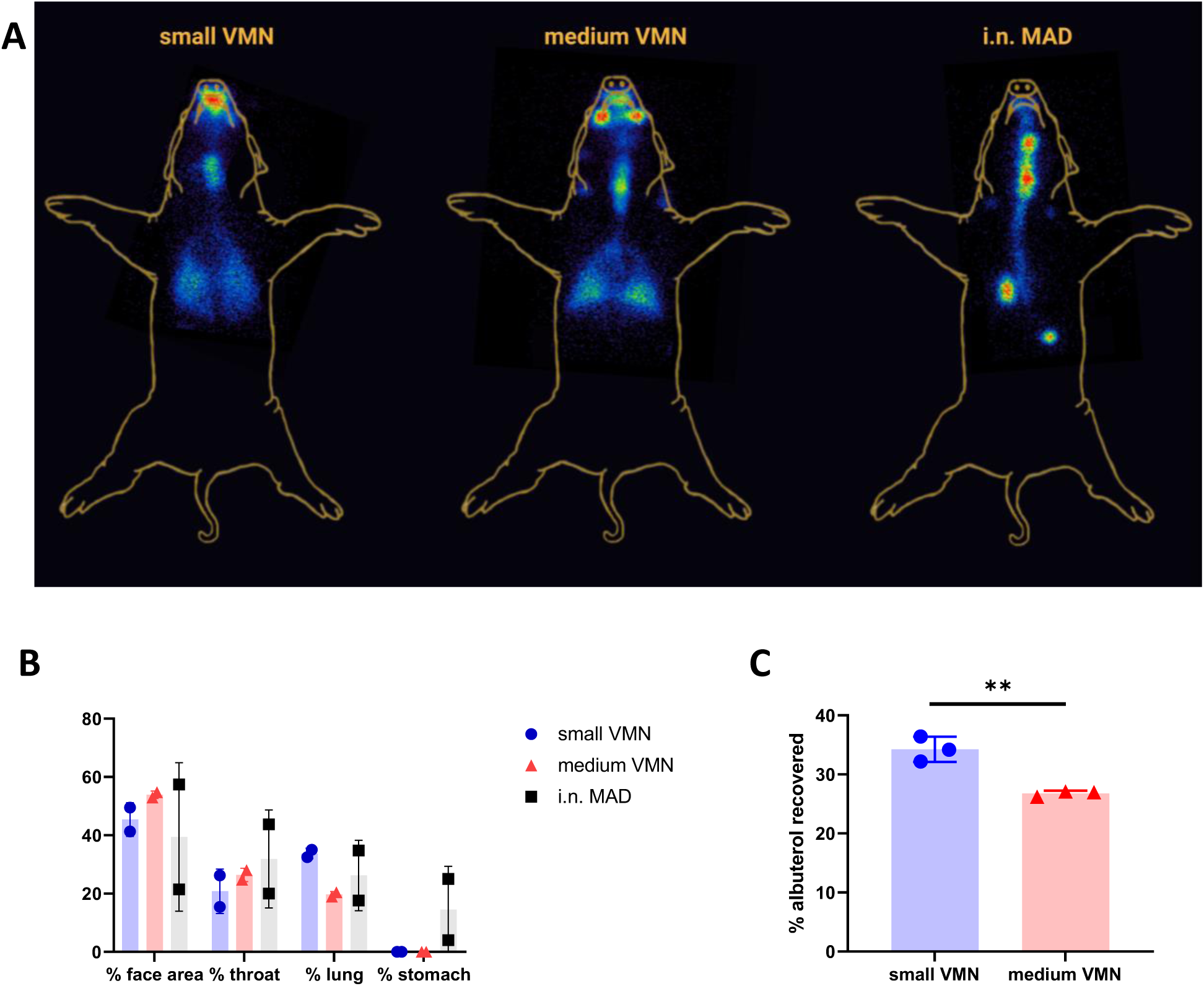
*In vivo* scintigraphy images and *in vitro* human head deposition. **(A)** Superimposed representative gamma counter images obtained using small or medium VMNs and i.n. MAD. Radioactive anatomical markers (visible as a circle in the two images) were positioned around the ears and thorax of the pig to enable *post hoc* image reconstruction. The arrow indicates the deposition in the stomach. **(B)** The reproducibility of scintigraphy was assessed by comparing the pilot study with the second leg where each animal had the same treatment as in the pilot. Counts in each region of interest for the two experiments are represented as % of total counts in the animal. **(C)** An adult human phantom was connected to medium and small VMNs using Ultra (Aerogen), a human aerosol mask, with 2 L/min oxygen flow. The phantom human head was connected to a respiratory pump using a lung/filter and the amount of albuterol quantified for three replicates for small and medium VMNs. Unpaired t-test was used for statistical analysis of albuterol deposition in the human phantom, using GraphPad Prism (8.3.0). Asterisks denote *p≤0.05, **p≤0.01, *** p≤0.001.

### Analysis of antibody and T cell immune responses

Neutralizing Ab titers, Virus specific IgG and IgA antibody titers and frequency of IFNγ secreting cells were performed as previously described (25). The phenotype of NP_217-225_ SLA tetramer binding specific CD8 T cells was established by flow cytometry on cryopreserved lymphocytes as described before (30). Briefly, biotinylated NP peptide loaded SLA monomers, kindly provided by Professor Andy Sewell, Cardiff University, were freshly assembled into tetramer with streptavidin BV421 (Biolegend, UK) and diluted with PBS to a final concentration of 0.1 μg/μl. Two million mononuclear cells were incubated with protease kinase inhibitor in PBS for 30 minutes at 37°C and then 0.3 µg of tetramer was added to the cells on ice for another 30 minutes. Surface staining with optimal antibody concentrations in FACS buffer (PBS supplemented with 2% FCS and 0.05% sodium azide) was performed on ice for 20 minutes (**Table II** for antibodies used). Samples were washed twice with FACS buffer and fixed in 1% paraformaldehyde before analysis using an LSRFortessa (BD Biosciences, UK). Data was analyzed by Boolean gating using FlowJo v10.6 (TreeStar, US). For intra-cellular cytokine staining, pH1N1 (MOI=1) was added to a 2 million BAL cells overnight at 37°C. Following a 5 hour incubation with GolgiPlug at 37°C, cells were stained with surface staining markers (**Table II**) before fixation and permeabilization using Cytofix Cytoperm (BD Biosciences, UK). Intracellular staining was then performed and the samples were analyzed using an LSRFortessa (BD Biosciences, UK).

**Table I.** Administered and recovered dose of ^99m^Tc-DTPA in pigs (mean ± SD)

| Device | Dose of radioactivity administered as % of dose loaded into device | Amount of radioactivity detected in the pig as % of dose administered | Amount of radioactivity detected in the pig as % of dose loaded into device |
| --- | --- | --- | --- |
| **Small VMN** | 49.2 ± 5.8 | 79.5 ± 23.3 | 37.8 ± 6.3 |
| **Medium VMN** | 52.4 ± 10.5 | 83.1 ± 14.0 | 42.0 ± 2.5 |
| **i.n. MAD** | 88.8 ± 0.5 | 64.3 ± 2.4 | 57.1 ± 2.4 |

**Table II.** List of antibodies used.

|  | **Antibody** | **Clone** | **Isotype** | **Fluorophore** | **Supplier** |
| --- | --- | --- | --- | --- | --- |
| **Tetramer**<br>**phenotyping** | Anti porcine CD3 | BB23-8E6-8C8 | mouse IgG2a | PE Cy7 | BD |
|  | Anti porcine CD4 | 74-12-4 | mouse IgG2b | PerCP Cy5.5 | BD |
|  | Anti porcine CD8 beta | PG164A | mouse IgG1 | PE | Biorad |
|  | Anti human CCR7 | 3D12 | rat IgG2a | BV711 | Biolegend |
|  | Anti porcine CD27 | B30C7 | mouse IgG1 | APC | Biorad |
| **ICS** | Anti porcine CD4 | 74-12-4 | mouse IgG2b | PerCP Cy5.5 | BD |
|  | Anti porcine CD8 beta | PG164A | mouse IgG1 | FITC | Biorad |
|  | Anti porcine IFN gamma | P2G10 | mouse IgG1 | AF647 | BD |
|  | Anti human TNF alpha | Mab11 | mouse IgG1 | BV421 | Biolegend |
|  | Anti porcine IL2 | A150D3F1 | mouse IgG2a | unconjugated | Invitrogen |
|  | Anti mouse IgG2a | RMG2a | N/A | PE Cy7 | Biolegend |

### Statistical analysis

Two-way ANOVA was used for statistical analysis of ROI data obtained in the *in vivo* scintigraphy study using GraphPad Prism (8.3.0), while an unpaired *t*-test was used for analysis of the pig and human breathing simulator as well as the *in vitro* particle size characterization experiments. Two-way ANOVA with Tukey post hoc test was chosen to compare the droplet size results of small and medium VMNs and i.n. MAD in the presence of S-FLU and ^99m^TC-DTPA. Kruskal-Wallis test was used to compare the NP_217-225_ CD8 population in the different tissues.

## Results

### *In vitro* characterization of droplets generated by VMN

An important component of targeted vaccine delivery to the RT is a device that can reliably and reproducibly deliver sufficient material to the site of interest. The site of vaccine deposition in the RT depends on, amongst other factors, the droplet size distribution of the inhaled aerosol, species specific anatomy and breathing pattern. Droplets larger than 10 μm predominantly deposit in the upper respiratory tract (URT) by inertial impaction, while droplets of less than 5 µm diameter are capable of reaching the lower respiratory tract (LRT); the trachea, bronchial and bronchiolar regions, as well as alveolar spaces (31, 32). Here we evaluated two VMNs producing different droplet sizes. Albuterol sulphate (salbutamol) a standard bronchodilator, was used as a tracer drug for aerosol characterization. Laser diffraction and cascade impaction analysis for the determination of aerosol droplet diameter, was performed in line with international regulatory submission standards. Analysis indicated that one VMN, termed “small VMN”, generated droplets with a volume mean diameter (VMD) of 3.5 µm and mass median aerodynamic diameter (MMAD) of 2.5 µm and the other, termed “medium VMN”, generated droplets with VMD of 4.5 µm and MMAD 3.5 µm (**Fig. 1A**).The small VMN generated significantly more droplets of less than 5 µm diameter, the fine particle fraction, than the medium VMN (**Fig. 1B**). These devices were selected with the expectation that the aerosols generated, would preferentially deposit in the lung, and minimise deposition within the URT.

In order to quantify the drug dose delivered to the LRT, we developed an *in vitro* model based on the breathing pattern and anatomical features of pig nasal cavities and trachea. A 3D printed pig head was attached to a breathing simulator with a collection filter, representing lung deposition, and albuterol was delivered using a custom-made mask. Because mammalian lung volume is influenced by bodyweight we modelled the breath of 15 kg (tidal volume of 115 ml) and 20 kg pigs (tidal volume of 150 ml), assuming 7.7 ml/kg bodyweight tidal volume (33).

These weights are in the range of the pigs used in the scintigraphy experiments described below. The small VMN deposited more albuterol on the filter (18.1% for 15 kg and 22.3% for 20 kg pigs) than the medium VMN (15.9% and 20.5% respectively) (**Fig. 1C**). These data showed that small and medium VMNs generated droplets with different size distributions which could potentially influence total lung deposition *in vivo*.

### Comparison of albuterol, technetium and influenza virus vaccine aerosol droplet size

To assess *in vivo* deposition of droplets, generated by the two VMNs or MAD in the pig we wished to conduct a gamma scintigraphy study with technetium (^99m^Tc) complexed with diethylenetriaminepentaacetic acid (DTPA)(^99m^Tc-DTPA), which is used as a radiotracer in clinical lung ventilation studies. Prior to using ^99m^Tc-DTPA as a surrogate for both albuterol and influenza virus particles such as S-FLU (34), we needed to establish that all three generated aerosols with similar droplet size. We used S-FLU because it is an attenuated influenza virus vaccine, identical in structure to wild type influenza but with a single replication cycle, making it safe for use in these experiments (27). The VMD of droplets produced by the two VMNs was comparable for albuterol, ^99m^Tc-DTPA, and S-FLU (differences of ± 0.4 µm are within the variability of the methodology) (**Fig. 1D**).

We also characterized the droplets generated using the MAD with ^99m^Tc-DTPA and S-FLU. The MAD consists of a syringe containing the formulation to which is attached an atomizer nozzle. Both solutions generated relatively large droplets (average VMD 86 µm). There was greater variability with the MAD (SD = 10 µm, CV = 9.9%) compared to the VMNs (SD = 0.05 µm, CV = 1.2%) (**Fig. 1E**).

The results indicate that *in vivo* deposition using ^99m^Tc-DTPA would be an accurate representation of the deposition of influenza virus or S-FLU vaccine particles. In addition, MAD generates a larger range of droplet sizes compared to the VMN nebulizers which might result in a different distribution in the porcine airways.

### *In vivo* deposition of ^99m^Tc-DTPA using VMNs and MAD

To evaluate RT deposition *in vivo*, ^99m^Tc-DTPA was administered to sedated pigs (n=3) using small and medium VMNs or i.n. with the MAD according to a randomised crossover experimental design (**Fig. 2A**). A 3- or 4-day interval between each administration of ^99m^Tc-DTPA ensured complete decay of the previous dose of radioactivity. Scintigraphic images of the sedated pigs were taken immediately after ^99m^Tc-DTPA administration using a gamma camera; this allowed the deposition of ^99m^Tc-DTPA in the animals to be visualized and quantified (as counts corrected for background and radioactive decay). The radioactivity associated with the various regions of interest (ROIs) was counted: face area (comprising ‘internal’ nasal cavities as well as ‘external’ facial skin), throat (trachea and oesophagus) and lung (**Fig. 2B**). Radioactivity detected below the lung area was considered to be in the stomach, as a result of swallowing nasal aspirate.

Images taken of each delivery system before and after administration to pigs revealed that the residual dose of ^99m^Tc-DTPA in the MAD was only 11% compared to approximately 50% with nebuliser plus mask; the latter was predominantly associated with the mask filter which is designed to capture exhaled air (**Table I** and **Fig. 2C**).The regional deposition of ^99m^Tc-DTPA in the pig was subsequently estimated as a percentage of the total amount detected (i.e. sum of counts associated with ROIs), as is standard practice. **Table I** indicates that a greater proportion of the administered dose could be accounted for in the pig with use of VMNs compared to MAD (79.5% and 83.1% versus 64.3%), equating to 37.8% to 57.1% of the dose actually loaded into the three devices.

It is well established that measurement of radioactivity using static 2-dimentional planar scintigraphy is not fully quantitative because it cannot reflect the 3-dimentional distribution of radiolabel in the body nor the dynamic nature of deposition and clearance and is influenced to varying degrees by scatter of radiation, tissue attenuation, distance from camera head, and by slight variations in background counts (35). Nevertheless, scintigraphy provides a useful evaluation of aerosol lung deposition.

Medium VMN deposited a significantly higher proportion (56%) of ^99m^Tc-DTPA in the face area compared to i.n. MAD administration (13.0%) (**Fig. 2D**). In the throat, i.n. MAD and small VMN deposited a slightly higher amount of ^99m^Tc-DTPA (30.4% and 35.9% respectively) than the medium VMN (18.7%). The i.n. MAD deposited a significantly higher proportion of the administered dose in the lung (51.0%) compared to 25.4% medium VMN while this difference did not reach significance for the small VMN (32.8%, p=0.16). In two out of three pigs the MAD delivered a proportion (7% and 9.8%) of the dose to the stomach. The droplet size distribution would normally be expected to prevent any deposition to the lungs using the MAD device but the finding of rapid drainage or aspiration to the lung and stomach with MAD is probably a consequence of the large volume used (1 ml per nostril) and perhaps the use of sedatives during administration.

The ‘face’ counts include those associated with deposition of ^99m^Tc-DTPA in the nasal cavities as well as those due to contamination of the external facial skin. We attempted to estimate the amount of facial skin contamination by removing external ^99m^Tc-DTPA with a skin wipe and imaging; revealing that more than 20% of the counts delivered by VMN to the face area were associated with the skin compared to 7% with the MAD (**Fig. 2E**). This fraction was unexpected, but could have arisen because of post-dose nasal drip or direct contamination of the external skin from the atomizer nozzle. Although wiping was not expected to recover 100% of the skin radioactivity, it nevertheless indicated significant deposition on the face when VMNs were used. In contrast, the majority of face counts appeared to be associated with the nasal cavities with the MAD (**Fig. 3A**).

Lung distribution patterns differed dramatically between VMNs and MAD. The MAD delivered a high dose of ^99m^Tc-DTPA to a relatively small area of the lung. In contrast, both nebulizers delivered droplets more uniformly throughout both lungs with small VMN resulting in higher and more consistent total lung deposition (mean ± SD 32.9% ± 1.6%) compared to the medium VMN (25.4% ± 6.8%) (**Fig. 2D & 3A**).

To establish the reproducibility of the scintigraphy technique, we compared the results between the pilot and second study legs, where the same devices were used in each animal. This comparison, although limited to 1 animal per device type, highlighted a potential for increased variability of i.n. MAD instillation compared to aerosol delivery by VMNs in all regions of the RT (**Fig. 3B**).

Together these data suggest that targeting to the pig lung is best achieved with use of a VMN. Although the MAD delivers a higher but variable proportion to the lung as well as the URT and the stomach, delivery via a nebulizer gives a much more even targeting of both lungs with deposition in the URT as well.

### *In vitro* deposition in a human head model

VMNs have been previously used for aerosol delivery of a measles vaccine in human (5, 10). To compare our findings with the human, we tested *in vitro* the delivery of albuterol to a human phantom, attached to a breathing simulator, using the VMN and Ultra facemask (Aerogen). A filter, connected to the distal end of the phantom, was also connected to the breathing simulator. Small VMN delivered significantly more albuterol (34.3%) to the ‘lung’ compared to medium (26.8%) (**Fig. 3C**). This comparison is in line with results in our *in vitro* pig model where small VMN reached the LRT more efficiently (**Fig. 1C**) and previous work in a human model (36). Similarly in humans, aerosol delivery using VMNs and a specific spacer (Aerogen Ultra) with mask deposited 34.1% in the LRT with 39.7% loss of the delivered dose, captured in the spacer device (37). The higher proportion of albuterol delivered to the human lung may be a reflection of the larger tidal volumes in humans compared to pigs as well as the more complicated anatomy of the pig URT.

### Intra-nasal delivery of influenza virus induces lung and systemic immune responses

Ferrets and mice are routinely challenged by nasal instillation with large volumes as are non-human primates for COVID-19 disease (38, 39). We routinely challenge pigs with influenza virus i.n. with MAD, but little information is available on the induction of lung immune responses after MAD infection. However, our deposition study highlighted the importance of evaluating the response in the lung where the majority of the dose is deposited using the MAD.

Because the MAD device targeted both the URT and LRT, we wished to assess immune responses in the entire RT generated by influenza virus, following MAD delivery. Four pigs were challenged i.n. using the MAD with pandemic swine H1N1 isolate, A/swine/England/1353/2009 (pH1N1) and culled 21 days post infection. We used inbred Babraham pigs, which have identical MHC (swine leucocyte antigen, SLA), allowing the assessment of dominant CD8 responses with peptide SLA-tetramers (30). Using *ex vivo* stimulation with pH1N1 T cell immune responses in tracheobronchial lymph nodes (TBLN), broncho-alveolar lavage (BAL) and blood were analyzed. Strong responses were induced in BAL (530 SFC per 10^6^ cells) and weaker responses in TBLN (105) and PBMC (166) (**Figs. 4A**). Intra cellular cytokine staining revealed that the majority of CD8 influenza specific T cells in the BAL secreted IFNγ and TNF (39.9%), followed by IFNγ only producers (35.7%) and 14.5% produced all three cytokines (IFNγ, TNF and IL-2), while CD4 T cells produced mainly IFNγ (49.3%) (**Fig. 4B**).

**Figure 4.**
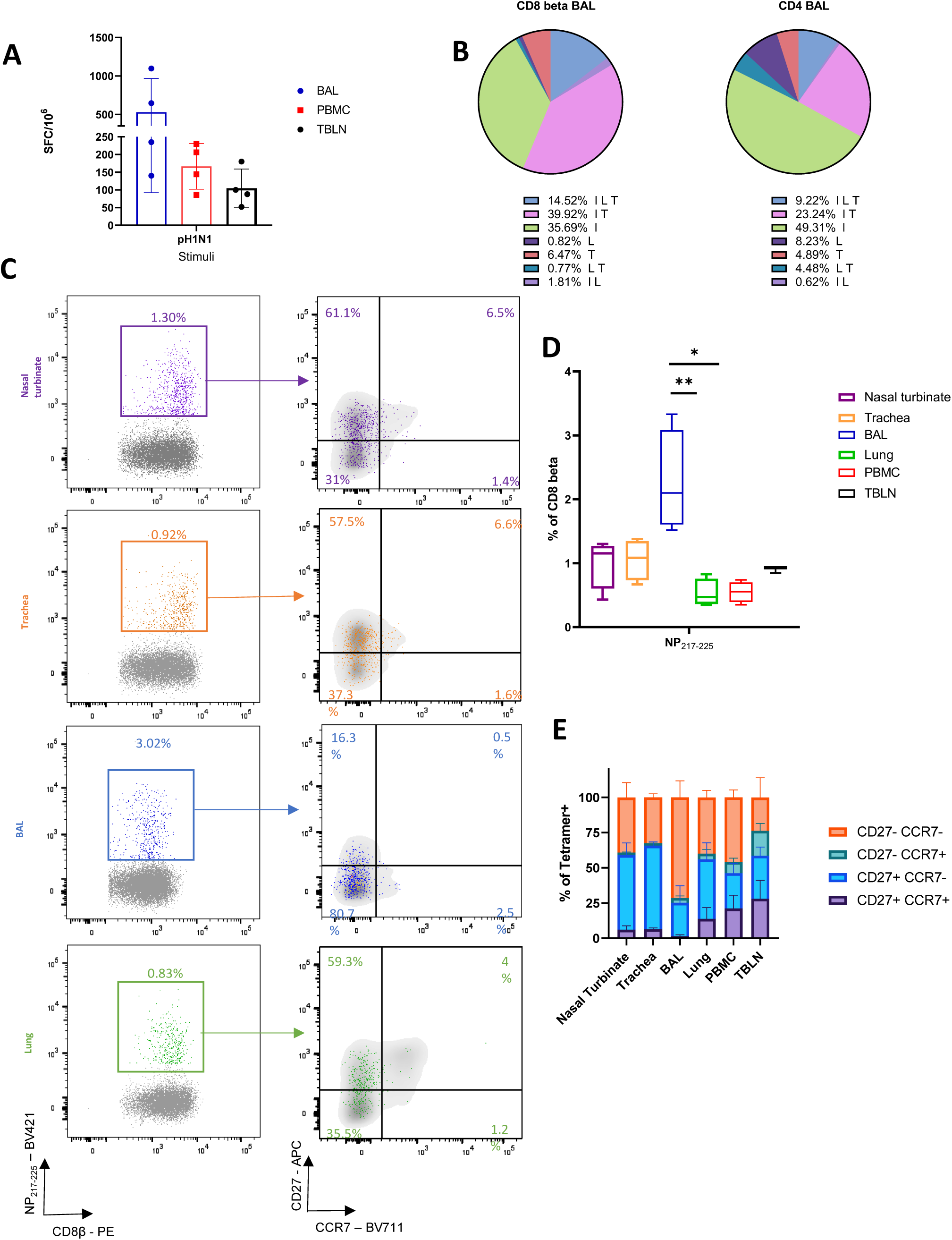
Cellular responses following i.n. MAD influenza challenge. Four Babraham pigs were challenged i.n. using MAD with pH1N1 and culled 21 days later. PBMC, nasal turbinates, trachea, lung broncho-alveolar lavage (BAL) and tracheobronchial lymph node (TBLN) were collected post mortem. **(A)** IFNγ secreting spot forming cells (SFC) in PBMC, BAL and TBLN were enumerated after stimulation with pH1N1. **(B)** BAL cells were stimulated with pH1N1 and cytokine secretion measured using intra-cytoplasmic staining. The pie chart shows the proportion of single, double and triple cytokine secreting CD8 T cells for IFNγ (I), TNF (T) and IL-2 (L) left and CD4 right. Percentages (mean of 4 pigs) are shown for each population. **(C)** Representative FACS plots showing the proportion of CD8+ NP_217-225_ cells in nasal turbinates (purple), trachea (orange), BAL (blue) and lung (green) expressing CCR7 and CD27. Tetramer+ CD8 T cells were identified with the following gating strategy: lymphocytes, singlets, alive (Zombie NIR™), CD3, CD8β and shown as NP_217-225_ vs CD8β. CCR7 and CD27 expression was analyzed in the CD8β (in gray) and compared to CD8+ NP_217-225_ (in colour). **(D)** Proportion of NP_217-225_ CD8 T cells in the respiratory tract and periphery. **(E)** Summary of CD27 and CCR7 expression in NP_217-225_ CD8 T cells in tissues. Kurskal-Wallis test was used to compare the NP_217-225_ CD8 population in the different tissues using GraphPad Prism (8.3.0). Asterisks indicates *p≤0.05 and **p≤0.01

We next enumerated and phenotyped the immunodominant NP_217-225_ tetramer+ CD8 T cells. BAL had the highest proportion of NP_217-225_ T cells (2.3%), followed by nasal turbinates (1%), trachea (1%) and TBLN (0.9%). Lung and PBMC had lower proportions (0.5%) (**Figs. 4C** and **D**). Our previous data using intravenous CD3 infusion indicates that more than 90% of BAL T cells are inaccessible to the blood and therefore are T_RM_ (25) and the present data indicated that 71.5% of the BAL cells had a highly differentiated CD27-CCR7-T effector memory (T_EM_) / T_RM_ like phenotype. In contrast the majority of tetramer positive cells in the nasal turbinates and trachea expressed CD27, but had downregulated CCR7, a less differentiated phenotype, while NP_217-225_ cells in the lung, TBLN and PBMC had more varied CD27 and CCR7 expression (**Figs. 4 C** and **E**).

We next assessed the humoral response, locally and systemically. A strong IgG and IgA ELISA response was detected in the serum and BAL with high neutralizing activity in serum (1:320 50% inhibition titre at day 21) and a much lower titre in BAL (1:15) (**Figs. 5A** and **B**). In the nose, IgA dominated over IgG and reached a plateau by day 14 (**Fig. 5C**). Taken together these results show that after i.n. MAD delivery a strong local T_RM_ response was induced in the BAL and URT, with a high titre of systemic and local Ab, although neutralizing Ab activity is low in the BAL.

**Figure 5.**
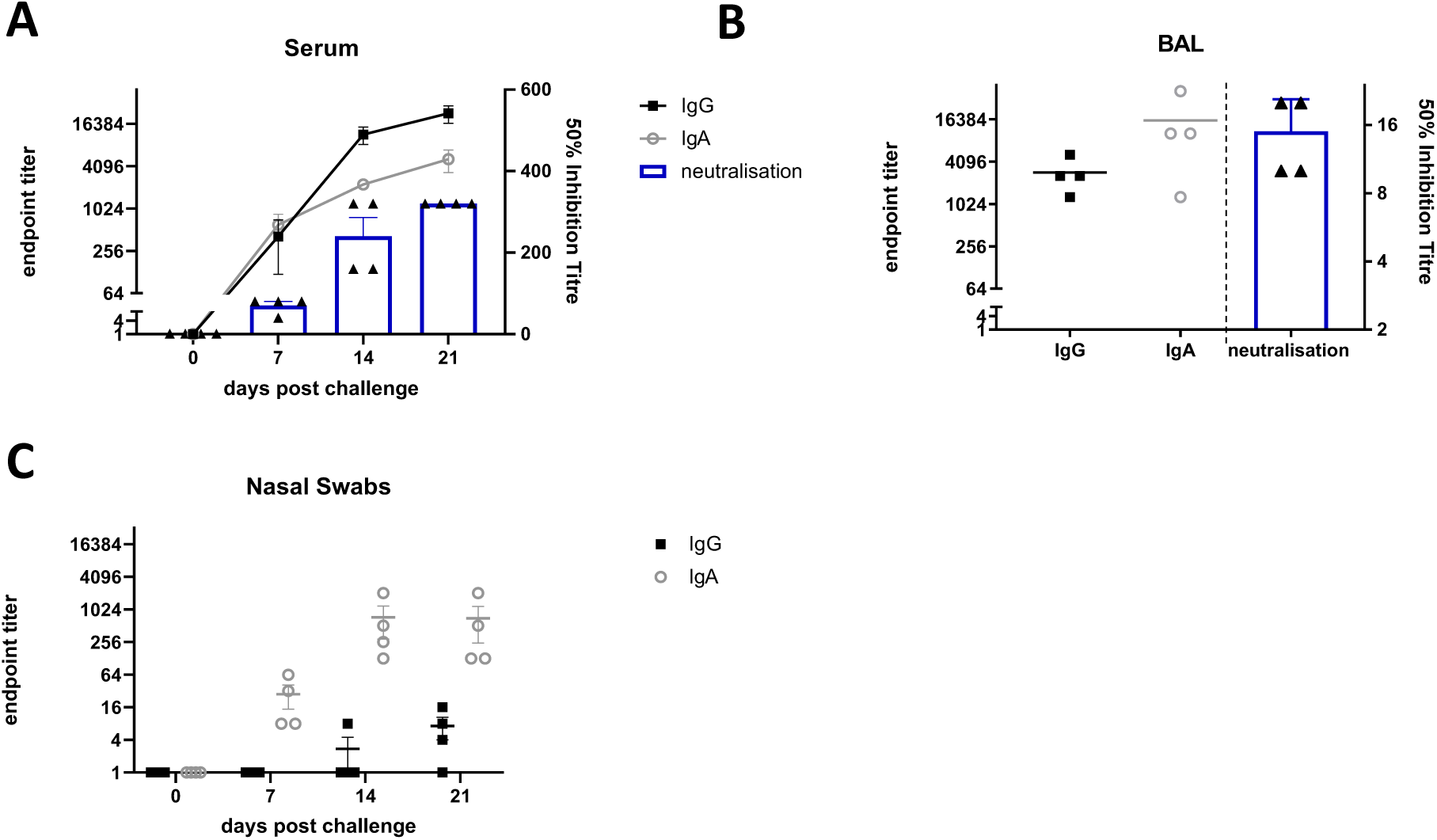
Antibody responses after i.n. MAD influenza challenge. Four Babraham pigs were challenged i.n. using MAD with pH1N1 and culled 21 days later. Serum samples were collected weekly IgA and IgG responses in serum **(A)**, BAL **(B)** and nasal swabs **(C)** were determined by ELISA and shown as dots. The mean 50% neutralization inhibition values in serum over time and BAL at 21 days post infection are shown in the columns.

## Discussion

In this study we evaluated the *in vitro* and *in vivo* deposition of droplets in the respiratory tract of pigs after i.n. MAD and VMN aerosol delivery. Surprisingly the MAD delivered the highest proportion of the dose to the lung but with very uneven distribution, localized in a relatively small area of one or both lobes. In contrast both VMNs delivered a lower proportion of the dose to the lung, but this was much more evenly distributed. In addition to the high lung deposition, some of the dose delivered with the MAD was found in the stomach. MAD delivery was much more variable both in terms of dose delivered to the lung, compared to very small variability with the nebulizers, and reproducibility within the same animal. The deposition pattern obtained with the MAD is most likely a consequence of the relatively large i.n. volume administered to the pigs.

Irrespective of the variability, the lung dose delivered by the MAD is large and therefore we asked whether concentration of influenza virus in a relatively small lung area affected the magnitude of immune responses. All MAD challenged animals generated strong immune responses and abundant local virus specific T cells in the BAL, that were multifunctional and resembled T_RM_ phenotypically. BAL NP217-225 CD8 T cells were mainly of T_EM_ phenotype as described in humans (40). For the first time we also characterized influenza specific trachea and nasal turbinate T cells in a large animal model. The proportion of tetramer specific cells in these sites was slightly lower than in BAL and their phenotype resembled CD8 T cells in human airways (40). Interestingly URT T cells in mice have been shown to be long lived and the less differentiated phenotype of pig URT T cells may indicate that these too turn over more slowly than the differentiated T_EM_ cells in BAL (41, 42). There was a smaller proportion of NP217-225 cells in the lung compared to BAL, probably due to the high vasculature of the organ and known difficulties in extracting T_RM_ from tissues (43). The phenotype of the NP217-225 cells in the lung was very similar to that in humans, CD27+/- CCR7-(44, 45).

Although a strong immune response was induced throughout the RT by MAD, only head to head comparison of the MAD and VMNs with the same virus or vaccine will determine unequivocally the best route to induce protective immune responses. Such a study will also determine if delivering large droplets is more or less effective in inducing protective immune responses and T_RM_ than aerosol administration. Studies with BCG given by aerosol or endobronchial instillation showed that the latter induced more T_RM_ (46). It will also be important to consider the volume to be delivered by the MAD. FluMist®, the commercially available LAIV, delivers 0.2 ml in total to humans intranasally, using the Accuspray device, which is very similar to the MAD, while in this study we delivered 1 ml per nostril, in an attempt to compensate for the larger swine nasal cavities. It is likely that a smaller volume will limit the deposition to the URT and therefore allow a better comparison between immune responses generated by delivery of a vaccine to the URT only, compared to delivery to the whole RT. Additionally, development of sophisticated VMNs that generates aerosol during the inhalation phase of the breath only, would minimise waste from exhalation, and maximise the lung dose. Such devices are currently used in humans, but not yet in pig models.

Restricting the response to the URT, by administering a smaller i.n volume, as in the case of LAIV, may not be optimal for lung protection, as studies in mice and ferrets suggest that induction of cross protective immunity against different types of influenza viruses is achieved most efficiently following vaccine delivery to the LRT (15, 27, 47). However, a barrier to delivering existing LAIV to the LRT is that LAIV retains some potential to replicate, raising safety concerns for lung delivery (48). Therefore, it remains to be formally tested whether lung targeting would make a difference to the protective efficacy of a vaccine directed against a respiratory pathogen. This might be of critical importance in situations with high exposure and risk of severe disease, such as medical staff in intensive care units. Although we have shown that administration of 2 × 1 ml by MAD reaches the LRT in sedated pigs, such volumes would be expected to be poorly tolerated in humans (49). Furthermore, the high variability of i.n. delivery raises the question over reproducibility of this strategy, although all four pigs infected by MAD produced strong immune responses.

In the field of respiratory medicine, the porcine model is becoming an increasingly important bridge between traditional small laboratory animal models and human medicine (21). It is important to take into account the more complex nasal anatomy of the pig compared to humans and that pigs are obligate nasal breathers, whereas humans may breathe through the nose and mouth. Nevertheless, the pig may be very useful in predicting responses to substances, including vaccines or infectious agents, administered to different regions of the RT.

It is not known whether it is important to target different regions of the RT to induce optimal protection against different RT infections. Using scintigraphy to establish the deposition of droplets by different delivery methods in parallel with vaccine administration, will allow the importance of different regions as inductive sites for immune responses, to be determined.

## Acknowledgements

We are grateful to the animal staff at the respective facilities for excellent animal care. We thank APHA for providing the challenge swine A/Sw/Eng/1353/09 influenza virus strain (DEFRA SwIV surveillance programme SW3401). We are grateful to Garry Bolton and Andrew Sewell, University of Cardiff for providing the SLA tetramers. We thank the Pirbright flow cytometry facility for their support.

## Author contributions

ET, VM, RM, AT, KB, MH conceived, designed and coordinated the study. VM, ET, AM, MH, EB, AT, MJ, RM performed animal experiments, processed samples and analyzed the data. ET, VM, PB wrote and revised the manuscript and figures.

